# Bacterial Cytochrome P450-catalyzed Post-translational Macrocyclization

**DOI:** 10.1101/2023.05.08.539676

**Authors:** Bei-Bei He, Zhuo Cheng, Jing Liu, Runze Liu, Zheng Zhong, Ying Gao, Hongyan Liu, Yong-Xin Li

## Abstract

Bacterial cytochrome P450s represent an emerging enzyme family that can modify ribosomally synthesized peptides to generate structurally complex macrocyclic skeletons. However, the functional sequence space of this type of enzyme is largely unexplored. In this study, we conduct a systematic genome mining of small ribosomal peptide-tailoring P450s from genomes of actinobacteria via a precursor-centric, primary sequence-, and structure-guided strategy. We uncovered 1,957 putative P450s, prioritized two representative families for functional study, and characterized two P450 enzymes that can respectively catalyze Tyrosine-to-Tryptophan and Tryptophan-to-Tryptophan crosslinks to form 3-mer or 4-mer macrocycle. These two P450 enzymes exhibit broad substrate selectivity, suggesting a promising starting template for engineering unnatural cyclic peptide construction. Our work expanded the enzymatic catalysis of P450s and could inspire the community to discover hidden peptide-modifying enzymes.

## Introduction

Ribosomally synthesized and post-translationally modified peptides (RiPPs), which are one of the most expansive families of natural products, possess a remarkable array of structural diversity and potential for bioactivity^1^. These peptides are found in various kingdoms of life, such as plants, fungi, and bacteria. They are biosynthesized using a similar mechanism whereby a ribosomal precursor peptide is modified by tailoring enzymes before being trimmed by a protease to release the mature product^1^. Cysteine thiols, serine and threonine hydroxyls, and aromatic residues, including tyrosine, phenylalanine, tryptophan, and histidine, tend to be activated by core tailoring enzymes to construct S-S, C-S, C-C, C-N and C-O bridges^2-7^. The RiPP families can be classified into lanthipeptides^2^, lasso peptides^5^, grasptides^8^, sactipeptides^9^, ranthipeptide^10^, thiopeptides^11^, cyanobactins^12^, and others, based on the presence of unique motifs introduced by distinct core tailoring enzymes^1^. These enzymes can be utilized in bioinformatic analysis to identify and classify biosynthetic gene clusters (BGCs) from sequenced microbial genomes via homology-based functional prediction, facilitating the discovery of novel natural products within known families.

However, identifying and classifying RiPP biosynthetic gene clusters (BGCs) that lack distinctive core enzymes, particularly for new RiPP families such as bacterial cytochrome P450-catalyzed RiPPs, remains a significant challenge. Due to the exceptional catalytic ability of P450s, distinguishing between RiPP-associated and non-RiPP-associated P450s can be difficult. The limited number of documented bacterial cytochrome P450-catalyzed RiPPs, including atropopeptides^6^, biarylitides^13^, and cittilins^14^ (Figure S1), typically contain one or more conserved aromatic residues such as tyrosine (Tyr, Y), tryptophan (Trp, W), or histidine (His, H). These conserved aromatic residues of core peptide allow P450 enzymes to activate substrate via homolysis of phenolic O−H bond or tryptophan indole N−H bond to form highly reactive radical intermediates^15^, which can undergo diverse coupling reactions to forge C-C, C-O, or C-N bonds. These oxidative couplings were also observed in cyclodipeptides (CDPs), such as diketopiperazine^16^, and non-ribosomal peptides (NRPs), including glycopeptide antibiotics vancomycin^17^ and lipopeptide arylomycin^18^, expanding the chemical space and complexity of macrocyclic peptides and highlighting the biocatalytic potential of P450 in peptide macrocyclization.

Significant efforts have been devoted to P450 discovery or engineering to establish a biocatalytic entry into these bioactive NPRs or CDPs^16, 18-23^. Such macrocyclization, however, either heavily limits to specific cyclodipeptides or requires a peptidyl carrier protein-tethered precursor for substrate recognition, which complicates the utilization of these P450 enzymes and limits the enzyme engineering. In contrast, ribosomal peptide-tailoring P450s with broad substrate selectivity provided a more straightforward biocatalytic route to these crosslinked cyclic peptides^24^. Identifying and characterizing new P450-catalyzed post-translational cyclization events are thus of significance for expanding the chemical diversity of strained macrocyclic peptides and enhancing our repertoire of biocatalytic tools. However, discovering P450-catalyzed RiPP BGCs is challenging due to the scarcity of characterized BGCs, extremely short precursor peptides, and the abundance of non-RiPPs-related P450 enzymes in bacterial genomes.

To address this problem, we first systematically analyzed aromatic residue-enriched small open reading frame (sORF)-P450 enzyme pairs in Actinobacteria using the SPECO workflow developed for RiPP BGC discovery^25^. Additionally, we introduced a new approach for uncovering P450-RiPP BGCs by combining multilayer sequence similarity network (MSSN) analysis of co-occurring sORF-P450 enzyme pairs and sORF-P450 complex prediction. This was realized by incorporating AlphaFold-multimer^26, 27^-based precursor-P450 complex screening into the genome mining workflow.

It allowed us to prioritize RiPP-associated P450s with high confidence. This approach enabled us to uncover 1,957 putative P450-catalyzed RiPP BGCs and prioritize 2 BGC families with unforeseen precursor patterns for functional study. Intensive BGC characterization and structural analysis uncovered two new P450 enzyme families, ScnB and MciB, which can respectively catalyze Tyr-Trp (C-C bond) and Trp-Trp (C-N bond) crosslinks within three or four residues. These widely distributed P450 enzymes in Actinobacteria represent a largely underexplored sequence space of peptide macrocyclization enzymes. Substrate scope investigation revealed that ScnB and MciB could accept non-native precursors, suggesting they could be promising starting templates for bioengineering to construct macrocycles.

## Results

### SPECO and AlphaFold-multimer-based genome mining reveals the landscape of P450-catalyzed RiPP BGCs

In recent years, advancements in DNA sequencing and bioinformatic algorithms for biosynthetic gene cluster (BGC) annotation have facilitated genome mining for discovering new compounds. However, identifying P450-catalyzed RiPP BGCs using current genome mining tools poses significant challenges compared to mining well-studied families. This is due to the short length of RiPP precursor peptides (e.g., the biarylitide precursor only contains five amino acids), a limited number of characterized BGCs, and the high abundance of P450 enzymes in bacterial genomes, especially in actinobacteria. To address this, we interrogated 21,911 actinobacteria genomes using SPECO workflow^25^ with an additional precursor filtering step: at least two aromatic residues (Trp, Tyr, or His) present in the C-terminal ten amino acids (Fig S2). This process generated 1,957 unique sORF-P450 enzyme pairs (dataset 1). To ensure the identification of P450-RiPP BGCs, we utilized a tailoring enzyme-precursor co-conservation approach based on primary sequence similarity. Enzyme-precursor multilayer sequence similarity network (MSSN) was constructed to find co-conserved sORF-P450 pairs (Figure 1A and 1B), including RiPP and non-RiPP sORF-P450 pairs (e.g., the ferredoxin-P450 system). To filter out these non-RiPP cases, we utilized the precursor-enzyme binding property in RiPP biosynthesis. To achieve this, AlphaFlod2-multimer was then implemented to calculate the sORF-P450 complex for BGC prioritization (Figure 1C).

**Figure 1.**
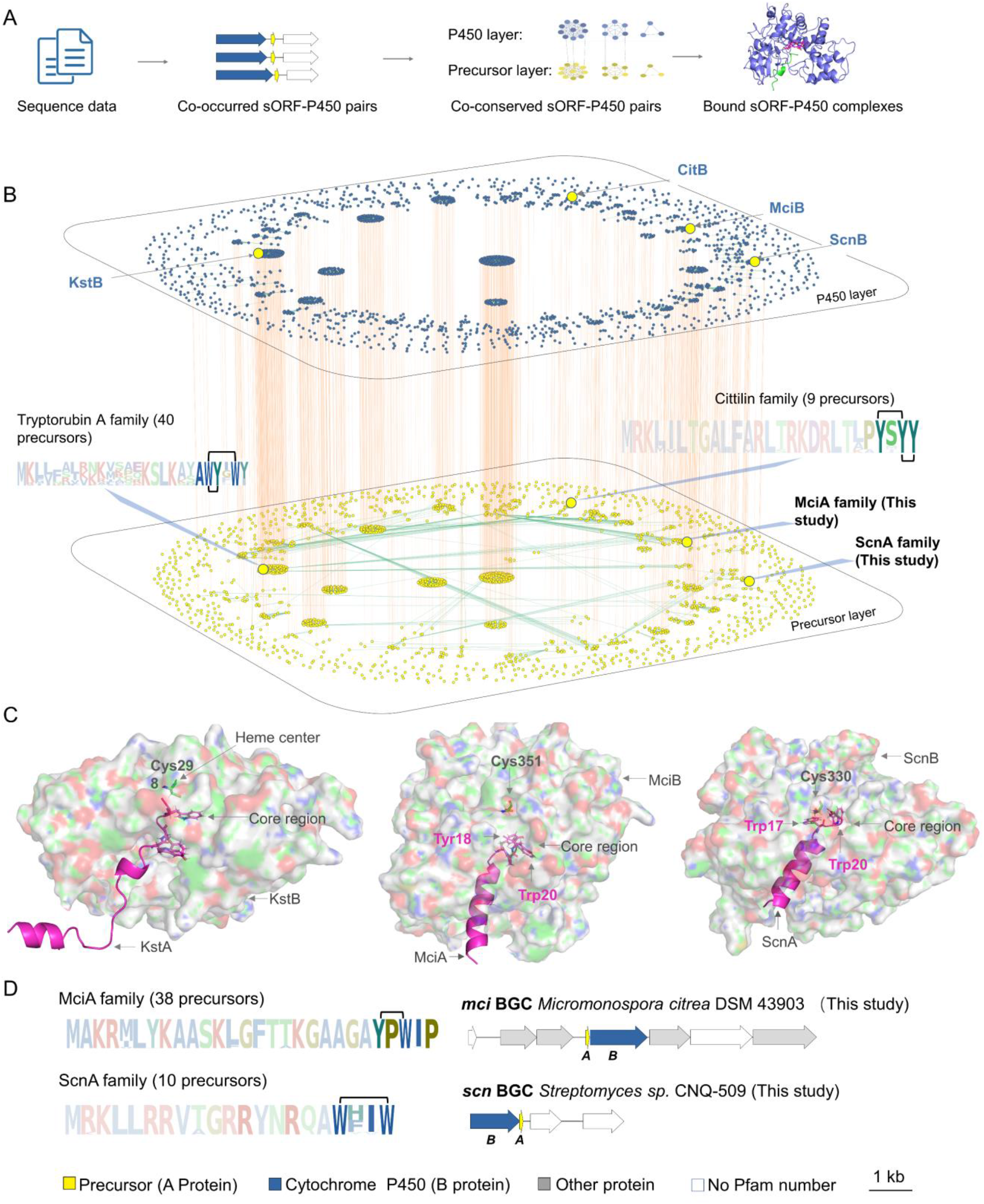
Genome mining of putative P450-catalyzed RiPP BGCs aided by SPECO and AlphaFold-multimer analysis. A. Genome mining workflow for discovering sORF-P450 pairs. B. MSSN analysis of 1,958 sORF-P450 pairs, with P450 enzyme and sORF sequence identities of 60% and 35%, respectively. C. Complex predictions of KstA-KstB, MciA-MciB, and ScnA-ScnB. D. Sequence logos and genomic contents of two BGCs characterized in this study.

Two families were prioritized based on (i) co-conservation of P450 and sORF, (ii) conserved C-terminal motif with aromatic residues, and (iii) predicted interaction in sORF-P450 complex and orientation of C-terminal core in P450 catalytic pocket. These BGCs include *mci* BGC family with conserved YPW motif and *scn* BGC family with W-X1-X2-W (X1 and X2 are variable residues) motif (Figure 1D). Two known families, including tryptorubin A and cittilin A were also covered in the MSSN (Figure 1B). Indeed, close analysis of the predicted tryptorubin A precursor-P450 complex revealed that the conserved Ser(-5) and Lys(-3) contribute multiple hydrogen bond interactions and Leu(-4) is extended into a highly hydrophobic pocket formed by multiple leucine residues (Leu253, 33, 255, 21, 257, 18, 333) from the P450 (Figure 1C and S3ABC). These putative precursor-P450-contacting residues are co-conserved in tryptorubin A BGC homologs (Figure S3DE), which may rationalize its biosynthesis.

We then chose one tryptorubin A BGC homolog, the *kst* BGC, from *Kitasatospora* sp. MMS16-CNU292 to further validate our approach. The precursor peptide hexa-histidine-SUMO-KstA alone and the hexa-histidine-SUMO-KstA-KstB system were both overexpressed in *E. coli* (Figure S4 and S5). Comparative ultra-performance liquid chromatography-high resolution mass spectrometry (UPLC-HRMS) analysis suggested that the peptide purified from KstAB coexpression system (G-KstA-4H, [M+5H]^5+^ = 606.9406) is 4 Da lighter than precursor (G-KstA, G-KstA, [M+5H]^5+^ = 605.9266) (Figure S5). HRMS analysis of the trypsin-digested fragment further confirmed that the 4 Da mass loss occurred in the C-terminal motif, consistent with the mass loss in tryptorubin A. These findings of co-conservation and interactions between P450 and sORF suggested the feasibility of our approach in prioritizing BGCs with high confidence.

As illustrated in Figure 1C, the predicted sORF-P450 complexes (KstA-KstB) of tryptorubin and those of Mci and Scn exhibit a similar arrangement, wherein sORFs are anchored by P450s in a manner that facilitates substrate activation through the heme center. The C-terminuses of KstA, MciA and ScnA are respectively buried in the catalytic pockets of P450s (Figure 1C). This orientation allows the ring-forming residues in KstA, MciA (Tyr18 and Trp20) and ScnA (Trp17 and Trp20) to fully extend to the heme center of the P450s (Figure 1C, S6-S7), which presumably provides a ready-to-activate conformation of the core peptide. The fully-buried binging pose of sORF distinguishes RiPP precursor-P450 pairs from other co-conserved but non-RiPP sequence pairs, e.g., the ferredoxin-P450 system has no such binding mode (Figure S8). Particularly, the MciA-MciB and ScnA-ScnB complexes revealed a handful of co-conserved hydrophobic residues in MciA (Phe9, Ala13-Ala17) and MciB (Leu17, 42, 306 and 284, Met307 and Trp74) that suggest a similar hydrophobic interaction between the precursor and P450 (Figure S6,S7). Furthermore, each predicted sORF exhibits an N-terminal α-helix and a C-terminal coil that may provide conformational freedom during ring construction (Figure S3B, S6B, and S7B). Taken together, the MciA-MciB and ScnA-ScnB sequence pairs meet all the requirements for the biosynthesis of RiPP, indicating the fidelity of these two BGCs. Our findings showcase the feasibility of using a combination of precursor-centric, primary sequence-, and structure-guided strategies to identify and prioritize rare RiPP BGCs.

### MciB catalyzes tyrosine-tryptophan crosslink

With prioritized BGCs in hand, we next explored the function of identified P450 enzymes. The *mci* BGC identified from *Micromonospora citrea* DSM 43903 was selected for characterization by a heterologous expression system using *E. coli*. Precursor gene *mciA* was cloned into multiple cloning site (MCS) 1 of plasmid pRSFDuet-1 with an N-terminal hexa-histidine-SUMO-TEV tag. P450 enzyme gene *mciB* was further inserted into MCS 2 site of pRSFDuet-1 to allow coexpression with tagged MciA in *E. coli*. Since the redox partner is essential for electron transfer in P450 catalytic cycle^28^, three exogenous redox partner pairs (spinach ferredoxin/ferredoxin reductase^22^, RhFRed^29^ and BM3Red^29^ were individually carried by pCDFDuet-1 vector and separately coexpressed with MciAB (Figure S4). After IPTG induction, Ni-affinity purification, and desalting, precursor peptides from MciA-only, MciAB coexpression system with or without exogenous redox partners were digested by tobacco etch virus (TEV) protease. Comparative UPLC-HRMS showed that precursor from MciA-only expression system (**1**, Gly-MciA, [M+5H]^5+^, *m/z* 514.6980) was 2 Da larger than that of purified from coexpression of MciAB (**2**, Gly-MciA*, [M+5H]^5+^, *m/z* 514.2949) without exogenous redox partner pairs (Figure 2A). Coexpression of redox partners with MciAB resulted in the same precursor modification as that observed in the sample without redox partners, indicating endogenous electron transfer system in *E. coli* is capable of modifying precursor. Tandem mass analysis of trypsin-digested fragments suggested a 2 Da mass loss in the YPW region (**3**, Figure 2B, S9), implying a crosslink may be formed between Tyr and Trp.

**Figure 2.**
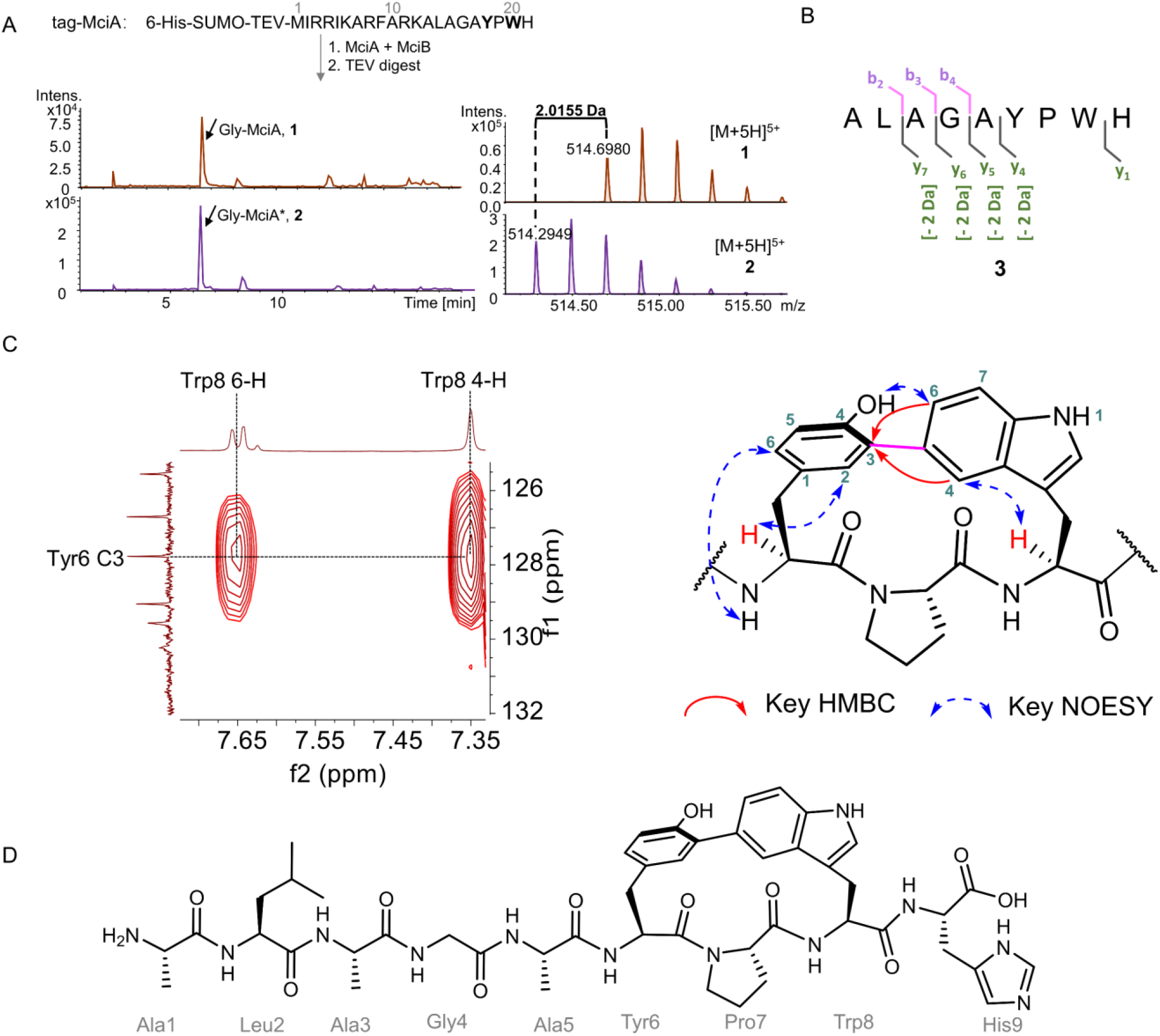
Characterization of *mci* BGC. A. UPLC-HRMS analysis of unmodified and modified MciA. B. Tandem mass analysis of modified fragment after trypsin digestion. C. A cutout of 600 MHz ^1^H-^13^C-HMBC NMR spectrum (left) and key HMBC and NOESY correlations (right) of **3**. D. Chemical structure of purified peptide **3**.

We then carried out a large-scale fermentation (10L), yielding 5 mg of **3** for 1D/2D nuclear magnetic resonance (NMR) analysis (Table S1, Figure S10-S19). Heteronuclear single quantum coherence (HSQC) and Heteronuclear multiple bond correlation (HMBC) spectra showed that H4 (7.36 ppm) of Trp8 (Trp 20 in precursor) is a singlet (Figure 2C), implying the adjacent C5 was substituted. H4 and H6 (7.65 ppm) of Trp8 showed HMBC corrections to a carbon with a chemical shift of 127.8 ppm, determined as C3 of Tyr6 (Figure 2C), suggesting a covalent bond was formed between these two residues. Together with the singlet signal of H2 (7.31 ppm) of Tyr6 and mass loss of 2 Da, confirmed the formation of a biaryl bond between Trp8 C5 and Tyr6 C3. This connection was further supported by the NOESY correction between H6 of Trp8 and the hydrogen atom of Tyr6 phenolic OH-group (9.14 ppm) (Figure 3C and S18). Advanced Marfey’s analysis of **3** using L/D-FDLA (5-fluoro-2,4-dinitrophenyl)-L/D-leucinamide) showed that alanine, proline, leucine and histidine residues were L-configured (Table S2). Based on the key NOESY corrections, Advanced Marfey’s results, and ribosomal origin of the precursor, we established that the formed cyclic peptide adopts a *S*_*a*_ axial chirality. To further confirm the axial chirality, we compared the experimental Electronic Circular Dichroism (ECD) spectrum of **3** with calculated isomers and found that the results supported the *S*_*a*_ axial chirality of **3** (Figure S20). We also used Gauge-Including Atomic Orbitals (GIAO) method to calculate the ^13^C chemical shifts of the proposed structure and found the results are in good agreement with experimental NMR data (Figure S21).

**Figure 3.**
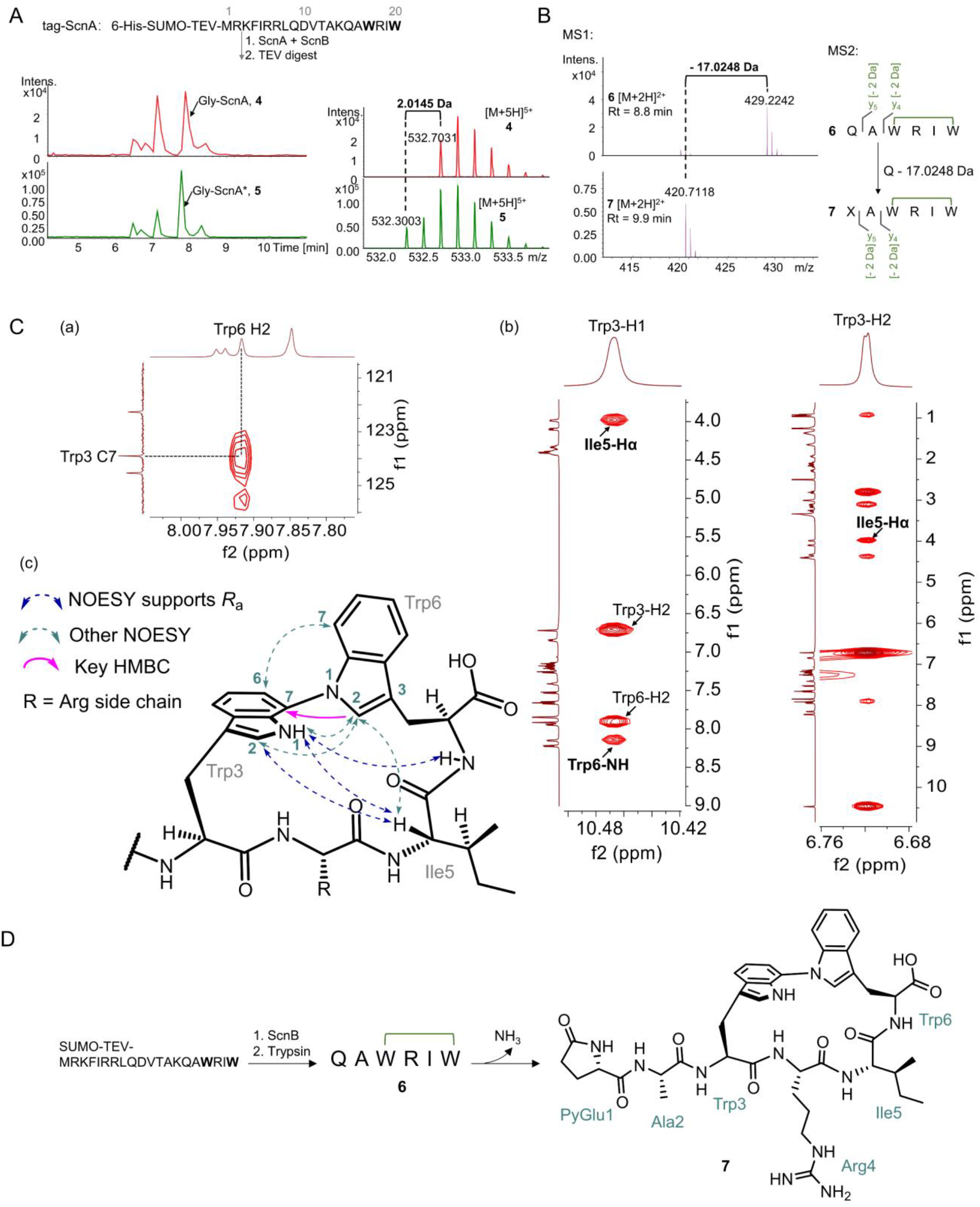
Characterization of *scn* BGC. A. UPLC-HRMS analysis of unmodified and modified ScnA. B. Tandem mass analysis of peptides **6** and **7**. The X residue in the C-terminus of **7** denotes Gln residue with 17 Da mass loss. C. Key ^1^H-^13^C-HMBC (a), NOESY (b) NMR spectrum, and core structure (c) of **7**. D. The structure and formation of **7**.

### ScnB catalyzes tryptophan-tryptophan C-N crosslink within 4-mer peptide

The *scn* BGC family features a conserved W-X1-X2-W motif in the C-terminus of the precursor, implying two Trp residues may be involved in modification catalyzed by the P450 enzymes. To verify our hypothesis, one BGC from *Streptomyces* sp. CNQ-509 was selected to explore potential enzymatic modification (Figure 1D). The precursor ScnA and P450 enzyme ScaB were cloned and overexpressed using the above-mentioned method. After purification and TEV digestion, comparative UPLC-HRMS analysis showed that peptide from ScnA-only (**4**, [M+5H]^5+^, *m/z* 532.7031) was 2 Da heavier than that of from ScnAB coexpression system (**5**, [M+5H]^5+^, *m/z* 532.3003) (Figure 3A). Tandem mass analysis of **4** and **5** suggested that mass loss occurred within the WRIW region (Figure 3B and S22), which was consistent with our hypothesis.

To release the modified WRIW region for NMR analysis, peptide **5** was digested by trypsin, yielding the desired fragment **6** ([M+2H]^2+^, *m/z* 429.2242) and a concomitant peak **7** ([M+2H]^2+^, *m/z* 420.7118) with 17 Da lighter than **6** (Figure 3B). Tandem mass analysis showed that **6** and **7** exhibited similar fragments and the same y4 and y5 ions, implying a mass loss of 17 Da occurred on the N-terminal Gln residue of **6** (Figure 3B and S22). Of note, a time-dependent decrease of **6** and an increase of **7** was observed (Figure S23), suggesting that the formation of **7** resulted from the spontaneous mass loss of **6**. We then purified peptide **5** from 20 L fermentation broth, followed by desalting and trypsin digestion. Since peptide **7** is more stable than **6**, we isolated 5 mg **7** for 1D/2D NMR analysis (Table S3 and Figure S24-S32). Only one characteristic indole NH (10.46 ppm) signal was observed in the proton NMR, implying that one of the two Trp residues may contribute an NH group for bond formation. The single peak of H2 (7.92, s) of Trp6 in proton NMR supported that its neighboring NH is substituted. The Trp6 H2 showed an HMBC correlation to a carbon with a chemical shift of 123.9 ppm, determined as C7 of Trp3 (Figure 3C-a). A C-N bond formation between C7 of Trp3 and N1 of Trp6 was established in the WRIW region, accompanied by a mass loss of 2 Da. This heteroatom bond formation led to a significant downfield shift of Trp3 C7 (123.9 ppm) compared to Trp6 (C7, 109.6 ppm, Table S3). The C-N bond also explains the absence of the proton on C7 of Trp3, as seen in the HSQC spectrum. Multiple NOESY correlations were observed (Figure 3C-bc and S32), including Trp3 H1 to Trp6 NH, Trp3 H1 to Ile5 Hα and Trp3-H2 to Ile Hα. Advanced Marfey’s analysis of **6** and **7** using L/D-FDLA showed that alanine, arginine, and isoleucine were L-configured (Table S4 and S5). The combination of key NOESY corrections, Advanced Marfey’s results, and ribosomal origin of the precursor provided evidence for *R*_*a*_ axial chirality in **7**. The ECD spectra of purified **7** exhibited a similar profile as the calculated data, further supporting the *R*_*a*_ axial chirality of **7** (Figure S33). Our calculations of the ^13^C chemical shifts for the proposed structure of **7** also agreed with the experimentally detected data (Figure S34). The N-terminus of peptide **7** features a pyroglutamate unit (Figure 3D), which was proposed to be formed via the cyclization of glutamine. This spontaneous cyclization of N-terminal glutamine was also observed in the *in vitro* reconstitution of KjaBURP for moroidin biosynthesis^30^. The pyroglutamate in **7** was converted into L-glutamate during acid hydrolysis (Table S5).

### MciB and ScnB enabled biosynthetic diversification of cyclic peptide

All the precursor families we characterized contain C-terminal Tyr or Trp residues, which may enable the P450 enzyme to activate the substrate by breaking the phenolic O−H bond or indole N−H bond. To assess the substrate tolerance of MciB and ScnB, we mutated the ring-forming residues to other aromatic residues, i.e., Tyr was replaced with His or Trp, and Trp was replaced with Tyr or His. (Table 1). For MciA mutants, HRMS analysis showed that W20Y single mutant was modified by MciB and yielded a monocyclic motif between the YPY region (Figure S35). No modification was detected in the remaining single or double mutations. ScnB, however, exhibited much higher substrate tolerance, as we found that positions 17 and 20 in ScnA could be Tyr or Trp. Both single (W17Y and W20Y) and double mutants (W17Y/W20Y) were modified by ScnB. The tandem mass of each modified fragment suggested the mass loss of 2 Da was introduced within the X1-R-I-X2 (X1 and X2 indicate Tyr or Trp) motif, which supported the formation of a 4-mer cyclic peptide (Figure S36-S38). Similar spontaneous deamination (-17 Da) was also observed in the trypsin-digested fragment (Figure S36-S37). No modification was observed in the shrunk (WRIW to WIW) or expanded (WRIW to WRGIW) core regions. The aromatic residue substitution assays demonstrated that both P450 enzymes MciB and ScnB exhibit broad substrate selectivity against key ring-forming residues. The variability of non-ring-forming residues, e.g., both hydrophobic (F18) and hydrophilic (H18) side chains, were observed in other ScnA homologs (Figure 1D), suggesting a diverse cyclic peptide library could be biosynthesized by these P450 enzymes.

**Table 1.**
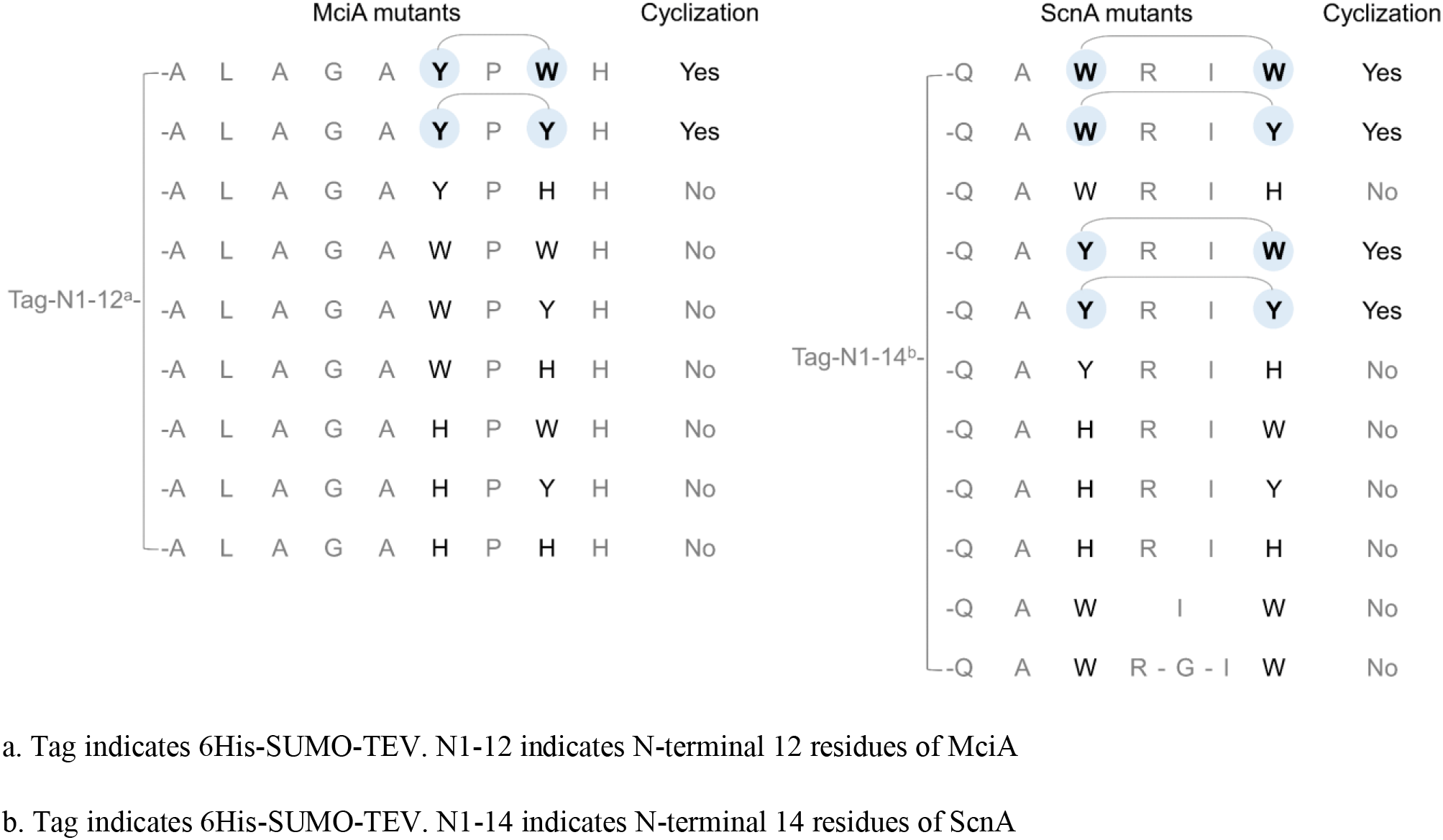
Cyclic peptide diversification assay of MciA (left panel) and ScnA mutants (right panel).

## Discussion

In this study, we adopted a systematic genome mining approach to identify small ribosomal peptide-tailoring P450s from Actinobacteria genomes. Our approach was precursor-centric, primary sequence-guided, and structure-guided, which allowed us to uncover previously unknown enzymology in RiPP biosynthesis. Despite the existence of over forty known RiPP families, our study highlights the vast potential for discovering new P450 enzymes by utilizing the fundamental biosynthetic logic of bacterial RiPPs. We identified two new families covering 48 precursor-P450 pairs, which we characterized by heterologous expression. Our findings also revealed the limitations of using homology-based strategies, such as BLAST, in discovering P450 enzymes, as all-to-all sequence alignment proved to be more challenging. For instance, all-to-all sequence alignment revealed that MciB and ScnB exhibit very low sequence identity (between 17.8% and 28.5%) with these known enzymes (Table S6), highlighting the challenges in discovering these P450 enzymes using homology-based strategy, e.g., using BLAST.

Despite the promising results of this study, it is important to note that we were unable to detect any natural products from the wildtype strains. This limitation suggests that further investigation is necessary to identify the factors or non-clustered genes that may affect the production of these natural products. Nevertheless, our study’s approach provides valuable insights into the discovery of new enzymology in RiPP biosynthesis and the potential for uncovering previously unknown P450 enzymes and their associated biosynthetic pathways. We believe that our workflow has significant implications for accelerating the discovery of untapped RiPP BGCs. While this manuscript was under preparation, Dong-Chan Oh and coworkers reported the cihunamides^31^, which resembles the crosslink formation in peptide **7**.

The discovery of new P450 enzymes in this study offers exciting opportunities for peptide engineering and the discovery of new natural products through a concise biosynthetic approach to preparing cyclic peptide libraries. While P450 enzymes found in NRP biosynthesis, such as glycopeptide, are effective at installing biaryl or diaryl ether crosslinks onto peptides, their utilization and engineering are complicated by the need for a peptidyl carrier protein-tethered precursor for substrate recognition^19^. In contrast, P450s involved in ribosomal peptide cyclization, such as ScnB, offer a more straightforward biocatalytic route to crosslinked cyclic peptides by catalyzing C-N bond formation between two Trp residues within a 4-mer peptide and exhibiting broad substrate selectivity. However, it is important to acknowledge that the lack of *in vitro* characterization of the identified P450 enzymes limits this study’s findings. While the comprehensive genome mining approach provides a valuable starting point for identifying untapped RiPP BGCs and new enzymology, further studies are needed to explore the catalytic efficiency of these enzymes fully. In vitro characterization of the P450 enzymes is necessary to fully understand their activity and potential applications, which could pave the way for peptide engineering and the discovery of new natural products.

Despite this limitation, our study presents a novel RiPP BGC genome mining workflow that combines primary sequence information and three-dimensional enzyme-peptide complex data to uncover two new P450 enzymes capable of modifying ribosomal peptides to form structurally complex molecules. This discovery highlights the remarkable catalytic versatility of P450s, and further investigation may uncover additional enzymes with unprecedented activity. Overall, this study’s findings offer promising opportunities for peptide engineering and the discovery of new natural products through a concise biosynthetic approach to preparing cyclic peptide libraries.

## Method

### General chemical reagents

All chemicals and culture medium, including kanamycin, streptomycin and isopropyl β-D-1-thiogalactopyranoside (IPTG) are purchased from either Sangon Biotech (Shanghai) or Sigma (USA). Marfey’s reagent Nα-(5-Fluoro-2,4-dinitrophenyl)-L-leucinamide (L-FDLA) and Nα-(5-Fluoro-2,4-dinitrophenyl)-D-leucinamide (D-FDLA) were purchased from TCI. Oligonucleotide primers were synthesized from BGI Genomics (Shenzhen). DNA polymerase Phanta Max Master Mix was purchased from Vazyme. TEV protease, endonucleases BamHI, Hind III, XhoI, NcoI, NdeI, XbaI, and Gibson assembly kit NEBuilder® HiFi DNA Assembly were purchased from New England Biolabs. Trypsin and PD-10 MidiTrap desalting columns were purchased from Sigma. HISPUR NI−NTA resin was purchased from Thermo Fisher. NMR solvent Dimethyl sulfoxide-d6 was purchased from Cambridge Isotope Laboratories.

### Bioinformatic analysis

SPECO pipeline was used to calculate co-occurred sORF-P450 pairs. Sequence clustering analysis of sORF and P450 enzyme sequences was conducted using mmseq2 with the following parameters: mmseqs --easy-cluster --min-seq-id [0.6 for P450, 0.35 for small peptide] --seq-id-mode 1 --cluster-mode 1 --similarity-type 2 --single-step-clustering --dbtype 1. Multilayer sequence similarity network (MSSN) analysis of sORF-P450 pairs were performed by employing Python package NetworkX and visualized by using Matplotlib.

### Strain preparation

All wild-type strains used in this study were purchased from the German Collection of Microorganisms and Cell Cultures GmbH (https://www.dsmz.de/). Cultivation was performed by following the guidance from DSMZ. *Escherichia coli* DH5α and *Escherichia coli* Rosetta (DE3) were separately used as cloning host and expression host.

### Cloning and plasmid construction

#### Redox partner gene synthesis and construction

For spinach ferredoxin and ferredoxin reductase, we synthesized the same nucleotide sequences (sequence id: *TRX-Spinach ferredoxin* and *Spinach ferredoxin reductase*) as reported by Prof. Xudong Qu, et al.^22^. The *E. coli* thioredoxin gene (highlighted as bold below) was fused to the N-terminal of ferredoxin to increase the solubility as reported by Prof. Xudong Qu, et al^22^. The maltose binding protein (MBP) was fused to the N-terminal of ferredoxin reductase to increase the solubility. These two fusion proteins were respectively cloned into multiple cloning sites 1 and 2 of pCDF-Duet-1 vector, generating pCDF-fdr-fdx.

#### Precursor and P450 enzyme cloning and plasmid construction

For precursor cloning, each precursor gene (*mciA, scnA*) was individually cloned by using each genome as a template and inserted into the multiple cloning site 1 of pRSF-Duet-1 with N-terminal His6-SUMO-TEV tag, yielding pRSF-SUMO-*mciA* and pRSF-SUMO-*scnA* plasmids. For precursor and P450 coexpression, corresponding P450 enzyme (*mciB, and scnB*) was respectively cloned and inserted into the multiple cloning site 2 of pRSF-SUMO-*mciA* and pRSF-SUMO-*scnA*, generating pRSF-SUMO-*mciAB* and pRSF-SUMO-*scnAB*.

### Transformation, protein overexpression and purification

Each redox partner, including pCDF-fdr-fdx, pCDF-Bre-BM3RED and pCDF-Bre-BM3RED, was individually transformed with *E. coli* Rosetta (DE3) to generate strain E. coli-redox. For precursor expression, plasmids pRSF-SUMO-*mciA* and pRSF-SUMO-*scnA* were respectively transformed with *E. coli* Rosetta (DE3). For co-expression of precursor and P450, plasmids pRSF-SUMO-*mciA* and pRSF-SUMO-*scnAB* were respectively transformed with *E. coli*-redox or *E. coli* Rosetta (DE3).

After transformation, a single colony of each recombinant strain was inoculated into 5 mL fresh LB broth with 34 μg/mL chloramphenicol (native resistance for *E. coli* Rosetta (DE3)) and 50 μg/mL kanamycin (for pRSF-Duet-1 backbone) and 100 μg/mL spectinomycin (when redox partner plasmids were needed) and shaken at 37 °C overnight. Then, 1 mL of the culture was incubated into 100 mL fresh TB medium (24 g/L yeast extract, 12 g/L casein hydrolysate, 9.4 g/L K_2_HPO_4_ and 2.2 g/L KH_2_PO_4_ and 8 ml/L glycerol) with antibiotics. After additional 2-4 hours of growth, the culture broth was cooled to 25 °C when OD _600_ reached 0.8. A final concentration of 0.5 mM IPTG was added to induce protein expression followed by a 20-hour induction at 25 °C, 120 rotations per minute (rpm.). The cells were harvested at 4 °C and centrifuged at 10,000 rpm for five minutes.

Each His-SUMO-tagged peptide was purified using a denaturing method. The following buffers were used for purification: denature buffer A (50 mM Tris, 500 mM NaCl, 6 M guanidine hydrochloride, adjusted pH to 8.0 using HCl), denature buffer B (30 mM imidazole in denature buffer A) and buffer C (50 mM Tris, 500 mM NaCl, 500 mM imidazole, adjusted pH to 8.0 using HCl). The cells were first resuspended using denaturing buffer A and then lysed using Sonics VCX-750 Vibra-Cell Ultrasonic Liquid Processor with 13 mm probe, 3 seconds on and 3 seconds off for 10 minutes. The cell lysis was then centrifuged at 4 °C, 10,000 rpm for 1 hour, and the supernatant was loaded into pre-equilibrated Ni−NTA resin (using denaturing buffer A) for three times to maximize target peptide yield. 5 mL denaturing buffer B was used to wash away non-specifically bound proteins. The target peptide was then eluted by using 1 mL buffer C for 4 times, i.e., 4 mL in total. The purified peptide was desalted using PD-10 desalting column. TEV protease and trypsin digestion were conducted under 30 °C for 1 hour and 37 °C for 24 hours, respectively. After digestion, the reaction mixture was centrifuged at 10000 rpm for 10 minutes, and the supernatant was subjected to UPLC-HRMS analysis.

### Mutation assay of precursors

MciA and ScnA mutants were constructed by using overlap PCR and confirmed by sequencing. Each mutant was individually transformed with *E. coli* Rosetta (DE3) for expression. Mutant purification, protease digestion and UPLC-HRMS analysis were conducted by following abovementioned methods.

### LC-HRMS analysis of purified peptides

For LC-MS analysis, Waters ACQUITY UPLC BEH C18, 130Å, 1.7 μm column was used. LC-HRMS experiments were performed on Thermo Scientific UltiMate 3000 UHPLC system coupled with Bruker impact Mass Spectrometer. The following LC method was used: column temperature (40°C), flow rate (0.2 mL/min), solvent A (0.1% formic acid in H_2_O), solvent B (0.1% formic acid in acetonitrile), 0-2 min (5% B), 2-15 min (5% to 95% B), 15-19 min (95% B), 19-21 min (5% B). The MS system was tuned using a sodium formate standard. All the samples were analyzed in positive polarity with m/z range from 150 to 1500 Th, using data-dependent acquisition mode.

### Semi-preparation of modified peptides

Larger scale fermentation of MciAB coexpression system was carried out in 20*500 mL TB medium. His-SUMO-tagged peptide was purified by using the same procedures mentioned above. To release target fragment (**3**), trypsin at a ratio of 1:100 (trypsin to peptide) was used to digest SUMO tag and linear part of the peptide in a 250 mL flask under gentle rotation (100 rpm) at 37 °C for 18 h. The trypsin -digested crude sample was then concentrated by a freeze dryer and redissolved into 5 mL methanol. The following LC method was used for peptide isolation: Column (semi-preparative Kinetx 5 μm XB-C18 100 Å, 250 * 10 mm), flow rate (3 mL/min), solvent A (0.05% TFA in H_2_O), solvent B (0.05% TFA in acetonitrile), 0-5 min (18% ACN), 5-40 min (18%-33% ACN), 40-45 min (100% ACN), 45-50 min (18% ACN). Purified **3** was freeze-dried, yielding about 5 mg white power. 250 μL Dimethyl sulfoxide-d6 was used to dissolve peptide **3** for NMR analysis.

Larger scale fermentation of ScnAB coexpression system was carried out in 30*500 mL TB medium. Purification, trypsin digestion and concentrating methods were the as mentioned above. For semi-preparation of peptide **7**, the following LC method was used: Column (semi-preparative Kinetx 5 μm XB-C18 100 Å, 250 * 10 mm), flow rate (3 mL/min), solvent A (0.05% TFA in H_2_O), solvent B (0.05% TFA in acetonitrile), 0-5 min (20% ACN), 5-40 min (20%-50% ACN), 40-45 min (100% ACN), 45-50 min (20% ACN). Purified **7** was freeze-dried, yielding about 5 mg white power. 250 μL Dimethyl sulfoxide-d6 was used to dissolve peptide **7** for NMR analysis. In parallel, peptide **6** was also purified at a small amount (less than 1 mg).

### NMR spectroscopy

^1^H NMR, ^13^C NMR, HSQC, ^1^H-^1^H COSY, TOCSY, and NOESY spectra were acquired on a Bruker Avance 600 MHz spectrometer with Cryoprobe, using DMSO-d6 as solvent. The Dimethyl sulfoxide-*d6* chemical shifts were used as the internal reference.

### Acid hydrolysis and Advanced Marfey’s analysis

For acid hydrolysis, dried 0.1 mg **3, 6** and **7** were individually prepared and dissolved in 1.5 mL 6M HCl in a glass vial and sealed tightly. Glass vials were then incubated in an oil bath under 110 °C for 16 hours. After hydrolysis, the aqueous phase was dried under the N_2_ stream. The remaining solid was dissolved in 100 μL H_2_O and equally transferred into two glass vials followed by adding 100 μL 1% L-FDLA and D-FDLA (dissolved in acetone), respectively. For each vial, 20 μL 1M NaHCO_3_ stock solution was added and incubated at 40 °C for 1 hour. After the mixture was cooled to room temperature, 10 μL 2M HCl was used to quench the reaction and diluted by adding 200 μL methanol. After centrifugation at 10000 rpm for 10 minutes, the reaction mixtures were analyzed by liquid chromatography low-resolution mass spectrometer.

### Electronic Circular Dichroism experiment

The purified peptides **3** and **7** were dissolved in H_2_O at the final concentration of 0.1 mg/mL. The CD spectrum of **3** and **7** were collected on a Jasco J-815 CD spectrometer with the following parameters: Band Width (5nm), Measure Range (400-180nm), Data Pitch (1.0 nm), Scanning Speed (100nm/min), D.I.T (2 sec).

For ECD calculation, the theoretical calculations were carried out using Gaussian 09^32^. At first, all conformers were optimized at PM6. Room-temperature equilibrium populations were calculated according to Boltzmann distribution law, based on which dominative conformers of the population over 1% were kept. The chosen conformers were further optimized at B3LYP/6-31G(d) in the gas phase. Vibrational frequency analysis confirmed the stable structures. ECD calculations were conducted at B3LYP/6-311G(d,p) level in H_2_O with IEFPCM model using Time-dependent Density functional theory (TD-DFT). Rotatory strengths for 30 excited states were calculated. The ECD spectrum was simulated using the ECD/UV analysis tool in Yinfo Cloud Computing Platform (https://cloud.yinfotek.com/) by overlapping Gaussian functions for each transition.

## Acknowledgements

This work is partially funded by the Hong Kong Branch of Southern Marine Science and Engineering Guangdong Laboratory (Guangzhou) (SMSEGL20SC02), the Research Grants Council of Hong Kong (27107320 and 17115322), and a Shenzhen Basic Research General Programme (JCYJ20210324122211031) to Y.-X.L. The authors would like to acknowledge and thank Yi-Man Eva FUNG, Jo Yip, Bonnie Yan and Dr. Yongqi Tian for their help in MS and NMR analysis.

